# ETV7 limits antiviral gene expression and control of SARS-CoV-2 and influenza viruses

**DOI:** 10.1101/851543

**Authors:** Heather M. Froggatt, Alfred T. Harding, Brook E. Heaton, Nicholas S. Heaton

## Abstract

The type I interferon (IFN) response is an important component of the innate immune response to viral infection. Precise control of interferon responses is critical; insufficient levels of interferon-stimulated genes (ISGs) can lead to a failure to restrict viral spread while excessive ISG activation can result in interferon-related pathologies. While both positive and negative regulatory factors control the magnitude and duration of IFN signaling, it is also appreciated that a number of ISGs regulate aspects of the interferon response themselves. However, the mechanisms underlying these ISG regulatory networks remain incompletely defined. In this study, we performed a CRISPR activation screen to identify new regulators of the type I IFN response. We identified ETS variant transcription factor 7 (ETV7), a strongly induced ISG, as a protein that acts as a negative regulator of the type I IFN response; however, ETV7 did not uniformly suppress ISG transcription. Instead, ETV7 preferentially targeted a subset of known antiviral ISGs. Further, we showed the subset of ETV7-modulated ISGs was particularly important for IFN-mediated control of some viruses including influenza viruses and SARS-CoV-2. Together, our data assign a function for ETV7 as an IFN response regulator and also identify ETV7 as a therapeutic target to increase innate responses and potentiate the efficacy of interferon-based antiviral therapies.

**One Sentence Summary:** ETV7 is an interferon-induced, repressive transcription factor that negatively regulates antiviral interferon-stimulated genes essential for controlling influenza virus and SARS-CoV-2 infections.

## Introduction

The type I interferon (IFN) response is a transient innate immune defense system that, upon activation by viral infection or therapeutic IFN treatment, induces the transcription of hundreds of interferon-stimulated genes (ISGs) (*1*). Many ISGs have characterized antiviral roles that restrict viral replication by either interfering with viral processes directly or altering cellular pathways required for viral replication (*2*). However, because replication mechanisms and points of interaction with the cell differ between viruses, individual ISGs have varying potencies against different viruses (*3*–*5*). As a result, unique combinations of ISGs are thought to mediate successful antiviral responses against distinct viruses (*1*, *6*).

The canonical activation pathway of the type I IFN signaling pathway induced by viral infection, or after therapeutic administration (*7*), is well understood (*8*, *9*). Extracellular IFN is bound by its cognate plasma membrane-localized receptor (IFNAR1/2). Downstream effectors (JAK proteins) are phosphorylated to then activate formation of the interferon-stimulated gene factor 3 (ISGF3) complex. Finally, the ISGF3 complex of STAT1, STAT2, and IRF9 translocates to the nucleus (*8*) and binds the interferon sensitive response element (ISRE), with the consensus DNA motif GAAANNGAAA, to activate transcription of ISGs (*10*).

As infection is cleared and virally-derived innate immune activators become scarce, interferon production is reduced and the interferon-stimulated gene response is downregulated. To facilitate this return to cell homeostasis, IFN induced negative regulators, such as SOCS1 (*11*) and USP18 (*12*), act at multiple levels in the signaling pathway (*13*). Thus, negative regulators of IFN responses are an important group of IFN-stimulated genes that control the duration of ISG induction and activity. Antagonism of interferon response negative regulators has been proposed as a mechanism to enhance host antiviral responses to clear infection, both alone and in conjunction with IFN treatment (*14*–*16*), and most recently during the COVID-19 pandemic (*17*). However, regulators working upstream of transcription impact ISGs indiscriminately, including suppression of the pro-inflammatory effectors, that when overabundant, can induce a cytokine storm; for example, agonists of an IFNAR-downregulating protein, S1PR1 (*18*), are proposed for use against pandemic influenza viruses (*19*) and SARS-CoV-2 (*20*) to instead limit excessive immune responses associated with interferon signaling. An ideal negative regulator of the IFN response for antiviral therapeutic targeting would enhance virus restricting ISGs specifically, without affecting pro-immune cytokines.

In addition to the upstream regulators that broadly activate or suppress IFN responses, there are interferon-induced transcriptional regulators that enhance, limit, or fine-tune ISG activity (*21*). Many ISGs themselves participate in innate immune signaling to amplify IFN and other pro-immune responses (*22*). Activators also add complexity by inducing non-canonical IFN response pathways or specific groups of ISGs. Interferon responsive factors (IRFs) 1 and 7 are ISGs and transcription factors that activate subsets of ISGs (*23*, *24*). Further, recent work has shown that ELF1 (E74-like ETS transcription factor) is induced by IFN and facilitates the expression of a group of genes not otherwise activated by the IFN response (*25*). Additional regulatory steps for ISGs, post-JAK/STAT signaling, likely exist to allow the cell to fine-tune its antiviral activity for an effective and appropriate response. While interferon-induced positive regulators of the IFN response are known to shape the complexity of ISG activation, reports of analogous roles for negative regulators remain conspicuously absent.

To address this gap in knowledge and identify genes able to shape the IFN response through negative regulation, we performed a CRISPR activation (CRISPRa) screen that selected for factors sufficient to prevent expression of an ISRE-containing IFN response reporter. We identified ETV7 (ETS variant transcription factor 7) as a negative regulator of the type I IFN response with a role in controlling the expression of specific ISGs. We further showed the ETV7-modulated ISGs are important for control of some IFN-sensitive respiratory viruses. Together, these data demonstrate ETV7 is a suppressive component of the complex ISG regulatory network that could be targeted to enhance specific antiviral responses against influenza viruses and SARS-CoV-2 (*1*, *26*).

## Results

### A CRISPR activation screen identifies ETV7 as a negative regulator of the type I IFN response

In order to identify negative regulators of the type I IFN response, we developed a type I IFN response reporter that included seven copies of the consensus interferon sensitive response element (ISRE) ahead of a minimal CMV promoter controlling expression of sfGFP (**Fig. 1A**). To make our reporter temporally specific, sfGFP was fused to a mouse ornithine decarboxylase (MODC) protein degradation domain to decrease its half-life (*27*). We stably introduced this construct into the A549 lung epithelial cell line along with a dCAS9-VP64 fusion protein and a MS2-p65-HSF1 activator complex required for the SAM CRISPR activation (CRISPRa) system (*28*). After clonal selection, 99.8% of the A549-SAM-IFN response cells expressed GFP in response to type I IFN treatment (**Fig. 1B and C**).

**Fig. 1.**
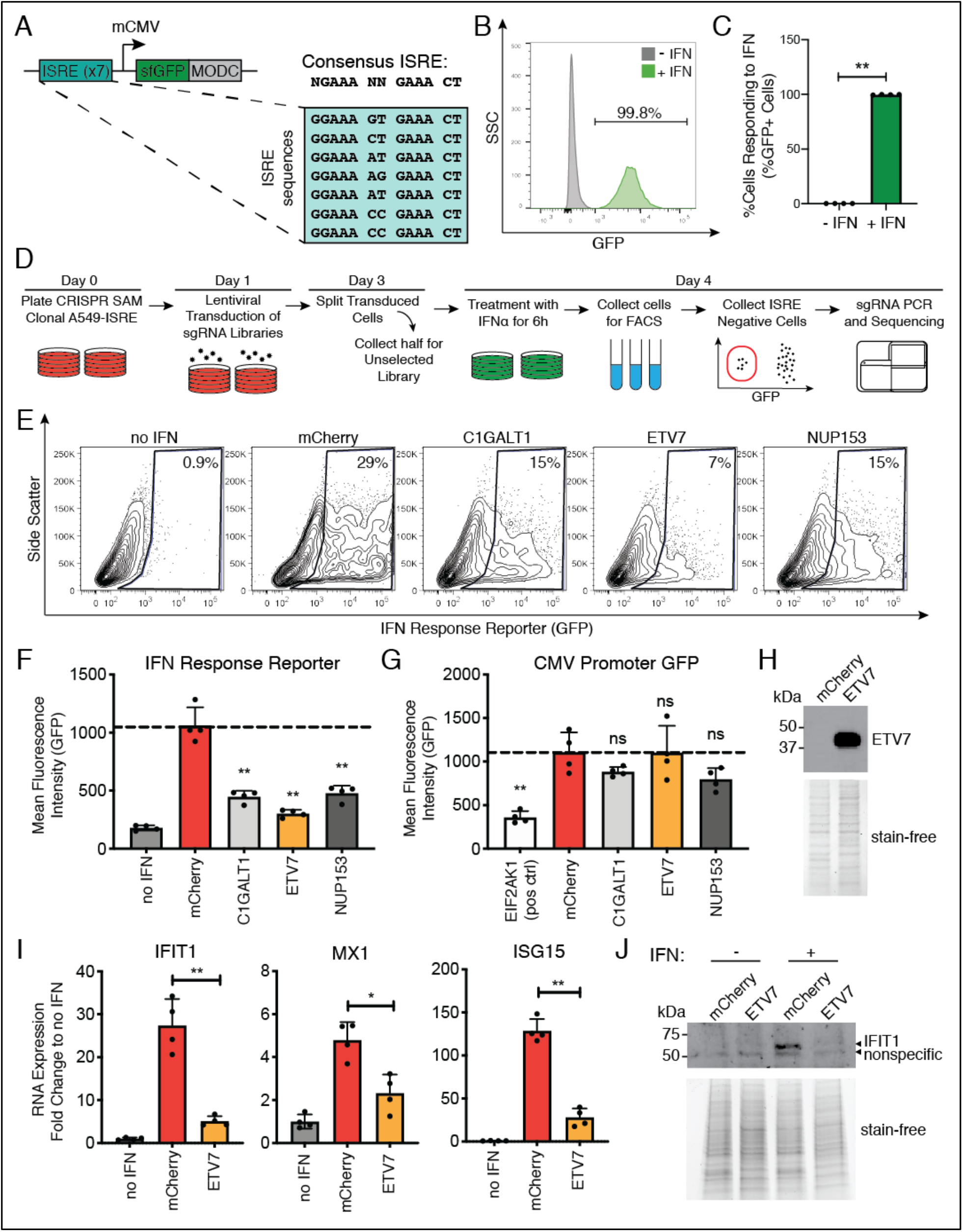
A CRISPR activation screen identifies ETV7 as a negative regulator of the type I interferon response. (**A**) Diagram of the IFN response reporter used to identify cells responding to IFN. ISRE = interferon sensitive response element, MODC = protein degradation domain. (**B**) Flow cytometry histogram and (**C**) bar graph of A549-SAM-IFN response cells before and after IFN-α treatment (1000 U/mL, 6 h) (data shown as mean ± SD, n=4, statistical analysis relative to untreated control). (**D**) Diagram of CRISPRa screen workflow to identify negative regulators of the type I IFN response. (**E**) Flow cytometry plots of 293T cells transfected with the IFN response reporter and overexpression plasmids for the indicated screen hits and then treated with IFN-α (100 U/mL, 9 h). (**F**) Quantification of **E** showing brightness of cells expressing GFP compared to the mCherry-expressing control (data shown as mean ± SD, n=4). (**G**) Brightness of cells expressing GFP from a constitutively expressing plasmid in cells overexpressing the indicated genes (positive control = EIF2AK1/HRI, shuts off translation) compared to control (data shown as mean ± SD, n=4). (**H**) Western blot showing ETV7 protein levels in 293T cells transfected with the ETV7 overexpression plasmid. Stain-free gel imaging was used to confirm equal loading. (**I**) Endogenous ISG mRNA expression levels measured using RT-qPCR after IFN-α treatment (100 U/mL, 9 h) (data shown as mean ± SD, n=4). (**J**) Western blot comparing IFIT1 protein levels in control and ETV7 overexpressing cells after IFN-α treatment (500 U/mL, 18 h). Stain-free gel imaging was used to confirm equal loading. For all panels: P-values calculated using unpaired, two-tailed Student’s t-tests (*p<0.05, **p<0.001) compared to mCherry-expressing control samples unless otherwise noted.

To perform the screen, we took the A549-SAM-IFN response cell line and introduced a lentivirus library containing sgRNAs designed to activate every putative ORF in the human genome (*28*) (**Fig. 1D**). After 48 hours, half of the cells were collected to determine the transduction efficiency and the remaining cells were re-plated for IFN stimulation. At 72 hours post-sgRNA introduction, the cells were treated with IFN-α and collected for fluorescence-activated cell sorting (FACS). During sorting, we eliminated reporter-positive cells and collected only cells that were nonresponsive to IFN as this population should theoretically be overexpressing a negative regulator of the IFN response. We performed two independent biological replicates of the screen and sequenced the sgRNA-containing amplicons derived from our input DNA, unselected transduced cells, and cells that were nonresponsive to type I IFN. Raw sequencing data was aligned, mapped, and subsequently analyzed using the MAGeCK pipeline (*29*) to generate z-score values for each gene. Genes were defined as “hits” if their z-scores exceeded two standard deviations from the mean, resulting in an overlap of 10 high-confidence genes between the two screen replicates (**Fig. S1A, Table S1, Data files S1 and S2**). We were seeking to identify regulators of the IFN response that are regulated by IFN themselves; therefore, we selected hits for validation previously reported to have at least a two-fold induction after IFN stimulation in the Interferome database (*30*). This analysis identified three hits (C1GALT1, ETV7, and NUP153) as potential negative regulators of the type I IFN response (**Table S1**).

To validate our three hits, and to eliminate any potential off-target effects of CRISPRa, we cloned the three ORFs and validated overexpression of the genes in 293T cells (**Fig. S1B**). Co-transfection of the overexpression plasmids and IFN response plasmid, followed by stimulation with IFN-α, resulted in lower reporter expression compared to a control mCherry-expressing plasmid (**Fig. 1E and F, Fig. S1C**). To verify this repressive activity was specific to the IFN response and the hits were not general inhibitors of transcription or translation, we transfected the overexpression plasmids along with a constitutively active GFP-expressing plasmid. We included a positive control (EIF2AK1/HRI), which is known to suppress host translation when overexpressed (*31*). None of the potential hits significantly reduced expression of the GFP-expressing plasmid (**Fig. 1G, Fig. S1D**). We therefore chose ETV7 for further characterization because: 1) NUP153 has previously been shown to control the distribution of STAT1 in the cell (*32*), 2) ETV7 had not been reported to play a role in the IFN response, and 3) it had the strongest inhibitory phenotype against the IFN response reporter.

After confirming overexpression of ETV7 at the protein level (**Fig. 1H**), we verified that the inhibitory effects of ETV7 were not restricted to the IFN response reporter plasmid. We collected mRNA and protein from IFN-α stimulated ETV7 overexpression cells to quantify effects on the expression of endogenous ISGs. ETV7 overexpression significantly repressed the induction of three prototypical ISGs (IFIT1, MX1, and ISG15) at the RNA level (**Fig. 1I**). The reduction of ISG expression during ETV7 overexpression was also demonstrated at the protein level for IFIT1 (**Fig. 1J**). These data together demonstrate that overexpression of ETV7 alone is sufficient to repress ISG induction by type I IFN.

### ETV7 acts as a transcription factor to repress IFN-induced expression

ETV7 is known to be a repressive transcription factor (*33*, *34*), although a role in repressing type I IFN responses has never been reported. To determine whether ETV7 acts as a transcription factor in this context, we generated a previously validated mutant of ETV7, called ETV7(KALK), which is unable to bind DNA (**Fig. 2A and B**) (*35*). Overexpression of ETV7(KALK) and stimulation with IFN-α had no measurable effect on expression of the IFN response reporter, in contrast to WT ETV7 overexpression (**Fig. 2C, Fig. S2A**).

**Fig. 2.**
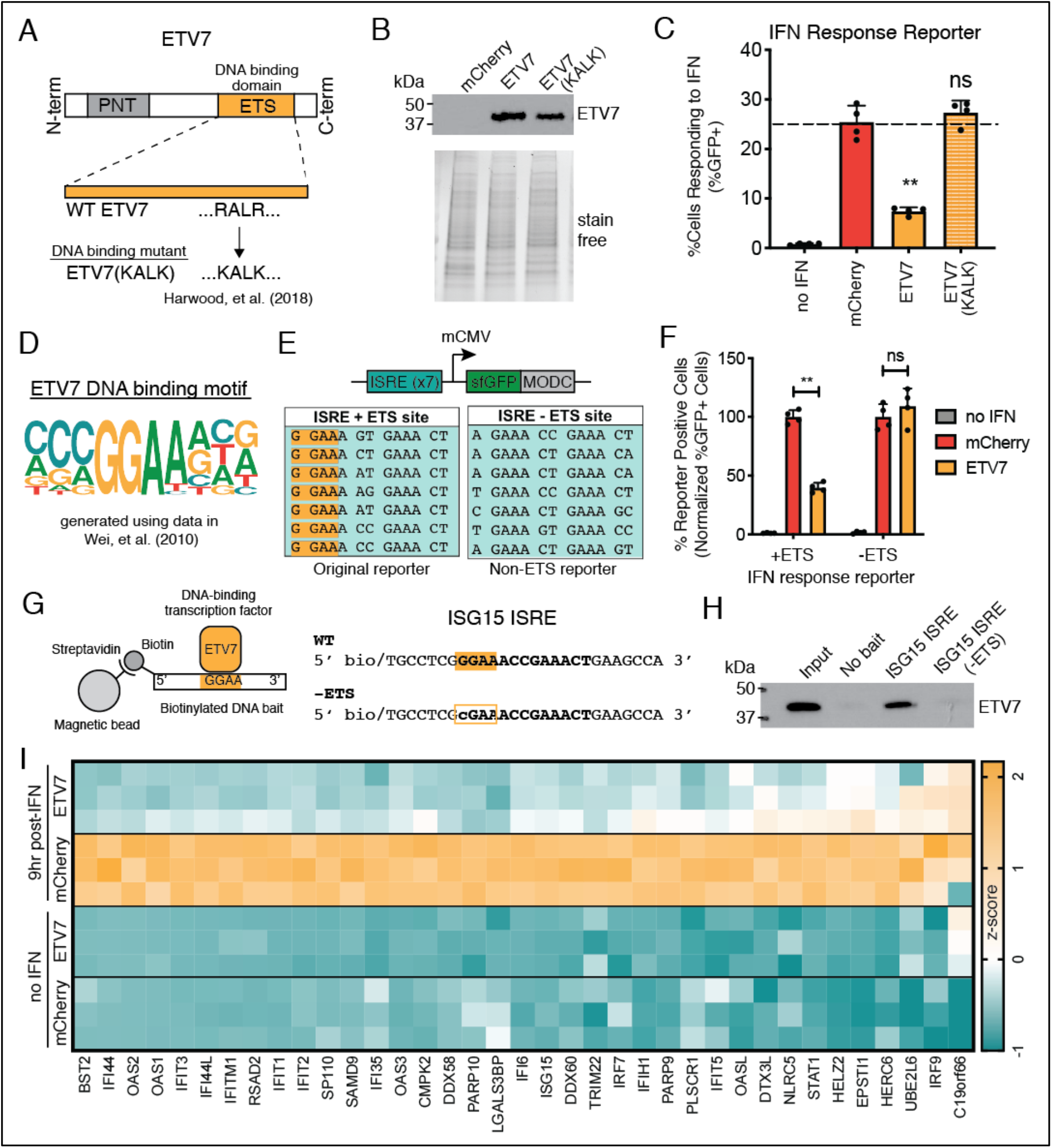
ETV7 acts as a transcription factor to negatively regulate the type I IFN response. (**A**) Diagram showing the ETV7 protein domains and amino acid changes made to generate the DNA binding mutant, ETV7(KALK). (**B**) Western blot showing ETV7 protein levels in 293T cells transfected with WT or DNA binding mutant (KALK) ETV7 expression plasmids. Stain-free gel imaging was used to confirm equal loading. (**C**) Percent of 293T cells expressing GFP from the IFN response reporter with overexpression of WT or DNA binding mutant (KALK) ETV7 after IFN-α treatment (100 U/mL, 9 h) compared to control (data shown as mean ± SD, n=4). (**D**) ETV7’s DNA binding position weighted matrix (PWM) generated using enoLOGOS (*99*) with data from Wei et al. (*36*). (**E**) Diagrams of the IFN response reporters containing (+ETS) and not containing (−ETS) potential ETV7 binding sites (ETS site, highlighted in yellow). (**F**) Normalized percent of 293T cells expressing GFP from IFN response reporters either containing or not containing ETS sites after overexpression of ETV7 and IFN-α treatment (100 U/mL, 6 h) compared to mCherry-expressing control (data shown as mean ± SD, n=4). (**G**) Diagram of the DNA pull-down experiment using biotinylated DNA and streptavidin-coated magnetic beads to show binding of a transcription factor to a specific DNA sequence. Sequences of biotinylated DNA bait used to show binding of ETV7 to the WT ISG15 ISRE (*93*) and loss of binding with mutation of the ETS binding site (−ETS). (**H**) Western blot of DNA pull-down using biotinylated oligos containing the ISG15 ISRE sequence in its wild-type form (WT) or with a single nucleotide mutation (−ETS) to eliminate the ETS site, incubated with nuclear lysate from cells expressing ETV7 with IFN treatment (100 U/ml, 9 h). (**I**) Heat map displaying RNA levels of genes upregulated at least 2.5-fold following IFN-α treatment (100 U/mL, 9 h) in control (mCherry) cells. Yellow = upregulated, blue = downregulated. For all panels: P-values calculated using unpaired, two-tailed Student’s t-tests (*p<0.05, **p<0.001) relative to IFN-stimulated, mCherry-expressing control samples.

ETV7 has been reported to bind the canonical ETS family DNA motif, GGAA (*36*), known as an “ETS” site (**Fig. 2D**). The original IFN response reporter design contained multiple ETS sites (**Fig. 2E**), which likely explains why it is negatively impacted by ETV7 in our screen. To test the requirement of ETS sites for ETV7 repressive activity against our reporter, we generated an IFN response reporter containing seven consensus ISREs from canonical ISGs that all lack ETS sites (ISRE −ETS) (**Fig. 2E**). We transfected the two reporter plasmids (ISRE +ETS and ISRE −ETS) independently into 293T cells and stimulated with IFN-α. As expected, both reporter plasmids responded to IFN treatment and were repressed by overexpression of known negative regulators of the type I IFN response (SOCS1, USP18) that function upstream of transcription (**Fig. S2B**). When ETV7 was transfected into cells with the IFN response reporters and stimulated with IFN however, the repressive activity of ETV7 was restricted to the reporter plasmid containing ETS motifs (**Fig. 2F, Fig. S2C**).

To provide evidence of ETV7 directly binding ISRE motifs containing ETS sites, we performed a DNA oligo-based pull-down experiment using the ISG15 ISRE sequence (**Fig. 2G**). We chose this ISRE because 1) it contains an ETS binding site, 2) it was included in our initial IFN response reporter, and 3) ETV7 was shown to impact ISG15 induction (**Fig. 1I**). In addition to the wild-type ISG15 ISRE biotinylated oligos, we tested biotinylated oligos with a single nucleotide mutation that eliminates the ETS site in the ISG15 ISRE (−ETS). We generated nuclear lysates from 293T cells overexpressing ETV7 and treated with IFN-α, incubated the lysate with either the biotinylated WT or −ETS ISG15 ISRE oligos, and pulled down the biotinylated DNA using streptavidin-coated magnetic beads. Western blotting for ETV7 protein revealed the transcription factor bound the WT ISG15 ISRE containing the ETS site, but not the mutated version of the promoter lacking the ETS site (**Fig. 2H**). Together, these experiments demonstrate that the repressive activity of ETV7 requires both its ability to bind DNA and the presence of ETS sites in the target ISG promoters.

### ETV7 non-uniformly represses IFN-stimulated gene expression

Since consensus ISREs can either contain or lack a GGAA motif (**Table S2**), we hypothesized ETV7 could differentially act on specific ISGs based on the presence or absence of ETS sites in their ISRE sequences and promoters. To perform an unbiased examination of the effect of ETV7 upregulation, we performed RNA sequencing in cells with or without ETV7 overexpression and IFN stimulation (**Data file S3 and Fig. S2E).** To identify the types of genes, processes, and functions impacted by overexpression of ETV7, we generated lists of the most downregulated genes during ETV7 overexpression, both before and after IFN treatment, and performed gene set enrichment analyses. This analysis technique takes a gene list and returns overrepresented terms from gene and protein databases (*37*). In the absence of IFN treatment, the most significantly enriched terms were not biological pathways, but rather ETS family transcription factor binding motifs (**Table S3**); this is expected, as ETV7 is an ETS family member and the entire family binds the same core motif, GGAA. In contrast, the genes specifically downregulated by ETV7 during IFN treatment were highly enriched for processes and pathways associated with responses to interferon and viral infection (**Table S4**). To visualize the effect of ETV7 upregulation specifically on ISGs, we generated a heat map of the genes induced at least 2.5-fold upon IFN treatment in our RNA sequencing experiment (**Fig. 2I**). Consistent with our hypothesis, we found some genes were suppressed by ETV7 overexpression more than others (**Fig. 2I**).

Although we had previously immunoprecipitated ETV7 with ISRE-containing oligos (**Fig. 2H**), we next wanted to demonstrate ETV7 occupancy on native ISG promoters for genes targeted by ETV7 via chromatin immune-precipitation followed by quantitative PCR (ChIP-qPCR). To enable these experiments, we first generated a FLAG-tagged ETV7 (**Fig. S2E**) and demonstrated that the tag does not disrupt its ability to repress the IFN response reporter (**Fig. S2F**). We next selected the promoter of IFI44L for ChIP analysis as it was both highly affected by ETV7 and also harbors many ETS sites within ISRE-like sequences in its promoter (**Fig. 2I and Fig. S2G and H**). To perform ChIP-qPCR, we generated sheared chromatin after cross-linking cells transfected with FLAG-tagged ETV7 and treated with IFN-α. We then performed immunoprecipitation using either nonspecific IgG or anti-FLAG antibodies raised in mice or rabbits. After qPCR analysis, we found no enrichment for a negative control region (the gene desert on chromosome 12) between the IgG and anti-FLAG samples (**Fig. S2I**). In contrast, DNA corresponding to the IFI44L promoter was significantly enriched with both the rabbit and mouse anti-FLAG samples compared to the nonspecific IgG samples (**Fig. S2I**). These experiments demonstrate that ETV7 acts as a transcription factor to bind and suppress ISG promoters that containing GGAA motifs.

### ETV7 loss enhances antiviral IFN-stimulated gene expression

Our experiments up to this point had primarily utilized overexpression of ETV7. One major disadvantage of this approach is the non-physiological kinetics and magnitude of ETV7 expression relative to what is observed after IFN stimulation in cells (**Fig. 3A**). To define the physiological effects of ETV7 induction during the IFN response, we performed a series of loss-of-function experiments. Our expectation was that the knockout of ETV7 would have the reciprocal effect on IFN responses as protein overexpression (*38*). We transduced A549-IFN response reporter cells (the original reporter with ISRE +ETS sites) with Cas9 and separately, two independent sgRNAs targeting ETV7, KO1(sg16667) and KO2(sg16668), generated pooled lines using antibiotic selection, and then stimulated with IFN-α. Both guides resulted in significantly more IFN-induced sfGFP expression compared to a control sgRNA (**Fig. 3B, Fig. S3A**).

**Fig. 3.**
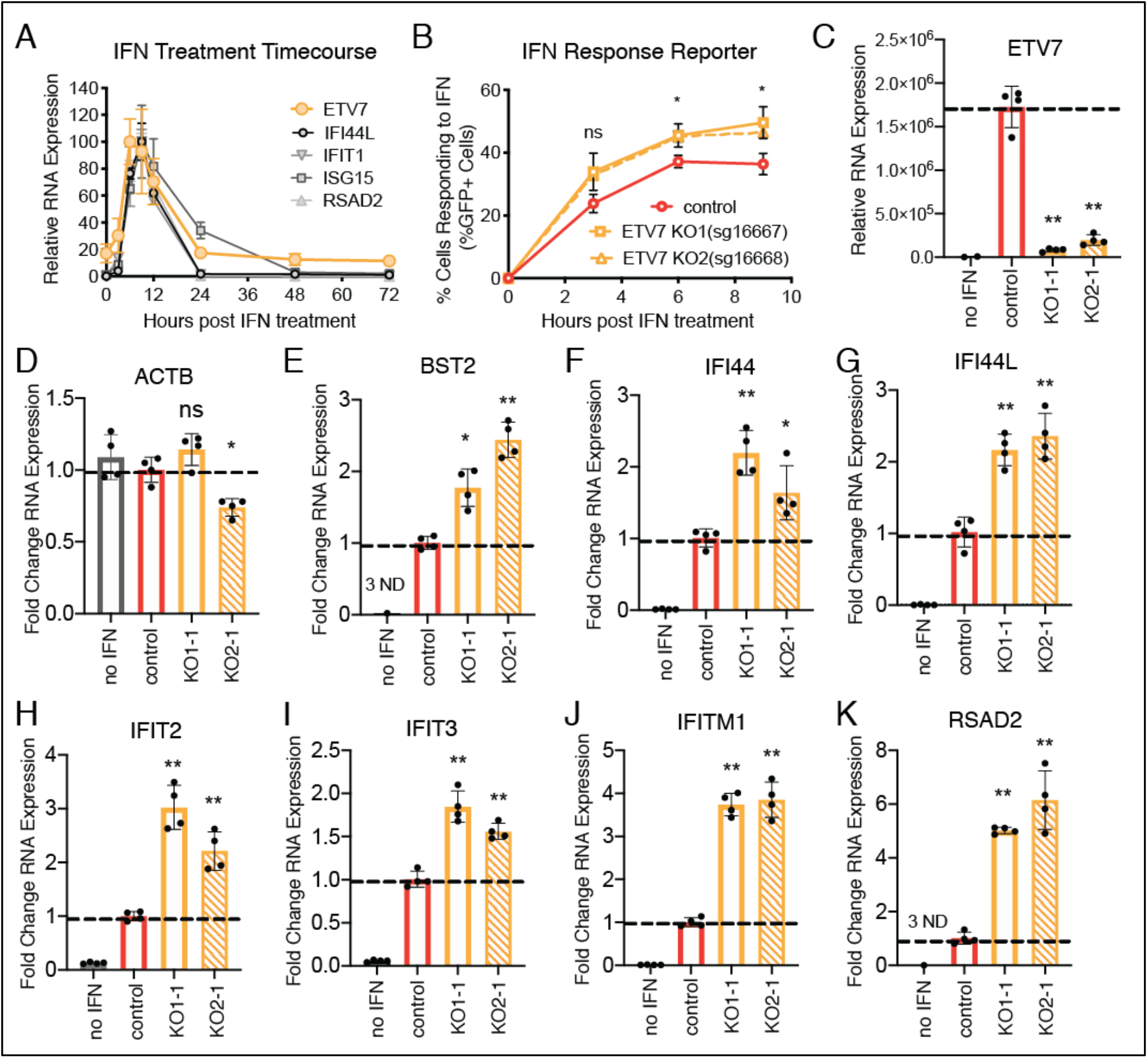
Loss of ETV7 enhances expression of ISGs. (**A**) ETV7 and other ISG mRNA levels in A549 lung epithelial cells after IFN-α treatment (100 U/mL, 6 h) (data shown as mean ± SD, n=4). (**B**) Percentage of cells expressing GFP from the IFN response reporter in two A549 ETV7 KO cell lines (pooled, 2 different guides) after IFN-α treatment (1000 U/mL, 6 h) compared to non-targeting control cells (data shown as mean ± SD, n=3). (**C**) mRNA levels of ETV7 in non-targeting control and ETV7 KO A549 clonal cell lines after IFN-α treatment (1000 U/mL, 6 h) (data shown as mean ± SD, n=4). (**D-K**) mRNA levels of a housekeeping gene (**D**) and ISGs (**E-***K*) in control and ETV7 KO A549 clonal cell lines after IFN-α treatment (100 U/mL, 9 h) (data shown as mean ± SD, n=4). For all panels: P-values calculated using unpaired, two-tailed Student’s t-tests (*p<0.05, **p<0.001) compared to IFN-stimulated, non-targeting sgRNA control samples.

We next generated clonal ETV7 knockout A549 lung epithelial cell lines (KO1-1, KO1-2) lacking the reporter and sequenced the resulting DNA lesions to confirm ETV7 knockout. Since ETV7 is normally only expressed after IFN stimulation, we treated with IFN-α and verified a reduction in ETV7 expression at the RNA level, presumably via nonsense mediated RNA decay (**Fig. 3C**). A representative housekeeping gene (ACTB) displayed no significant increases in transcription in any of the ETV7 KO clones (**Fig. 3D**). We then selected twelve ISGs (IFITM1, RSAD2, OAS1-3, IFIT1-3, IFI44L, ISG15, BST2, and IFI44) that were highly impacted by ETV7 in our RNA sequencing results for RT-qPCR analysis after IFN treatment. Compared to clonal lines containing a non-targeting guide, the ETV7 KO lines showed significant increases in induction of each of the twelve ISGs (**Fig. 3E-K, Fig. S3B-F**). Thus, the physiological induction of ETV7 after IFN stimulation affects the expression of ISGs.

### Suppression of ETV7 enhances IFN-mediated control of influenza viruses and SARS-CoV-2

In trying to predict the physiological significance of excessive induction of these ETV7-regulated ISGs during the type I IFN response, we noted that many have recognized antiviral functions (*2*, *39*, *40*). To determine whether the effects of ETV7 suppression of ISG expression were relevant in the context of a viral infection, we wanted to identify a virus restricted by the ISGs most affected by ETV7 (*41*). Many of these genes with well-recognized antiviral functions (IFITM1, IFIT1-3, OAS1-3, BST2, RSAD2) have been reported to play important roles in the restriction of influenza viruses (*42*). IFITM1 has been shown to prevent viral entry (*43*), OAS proteins activate RNase L to degrade viral RNA (*44*), IFITs bind viral RNA and promote antiviral signaling (*45*), and BST2/Tetherin and RSAD2/Viperin restrict viral budding and egress (*46*, *47*). Therefore, we hypothesized that ETV7 dysregulation would affect influenza virus replication and spread.

We first infected our ETV7 KO A549 cells with a laboratory-adapted H1N1 influenza A virus (IAV), A/Puerto Rico/8/1934 (PR8). Using a hemagglutination (HA) assay to measure the number of viral particles released over time, we observed reduced virus production in our ETV7 KO cells compared to control cells (**Fig. 4A**). This was the anticipated outcome because loss of a negative regulator (i.e. ETV7) is expected to enhance expression of antiviral ISGs. We also measured infectious viral titers and found a significant reduction in our ETV7 KO cells compared to control cells (**Fig. S4A**). Using a fluorescent reporter strain of PR8 (PR8-mNeon) (*48*), we next visualized infection and spread. As expected, we observed fewer cells expressing mNeon in ETV7 KO cells using both microscopy (**Fig. 4B**) and flow cytometry readouts (**Fig. 4C**). We also tested whether this phenotype would extend to a more contemporary H1N1 IAV strain, A/California/07/2009 (Cal/09), as well as an unrelated Victoria lineage influenza B virus strain, B/Malaysia/2506/2004 (Mal/04) (*49*). Using fluorescent reporter strains of these viruses, we observed significant decreases in the number of Cal/09- and Mal/04-infected cells when comparing ETV7 KO cells to control cells (**Fig. 4D and E**). In order to rule out non-IFN-related effects of ETV7 on inhibition of influenza viral replication, we also performed these experiments with a non-influenza virus. We selected Sendai virus (SeV) because, although it is known to induce a strong IFN response, previous reports (*50*) show SeV replication is relatively unaffected by type I IFN. Furthermore, many of the ISGs most impacted by ETV7 have little to no effect against SeV, including IFITM1 (*51*), IFIT1 (*52*), and BST2/Tetherin (*53*). Using of a fluorescent reporter SeV, we observed no significant change in the number of infected cells when comparing ETV7 KO cells to control cells (**Fig. 4F**).

**Fig. 4.**
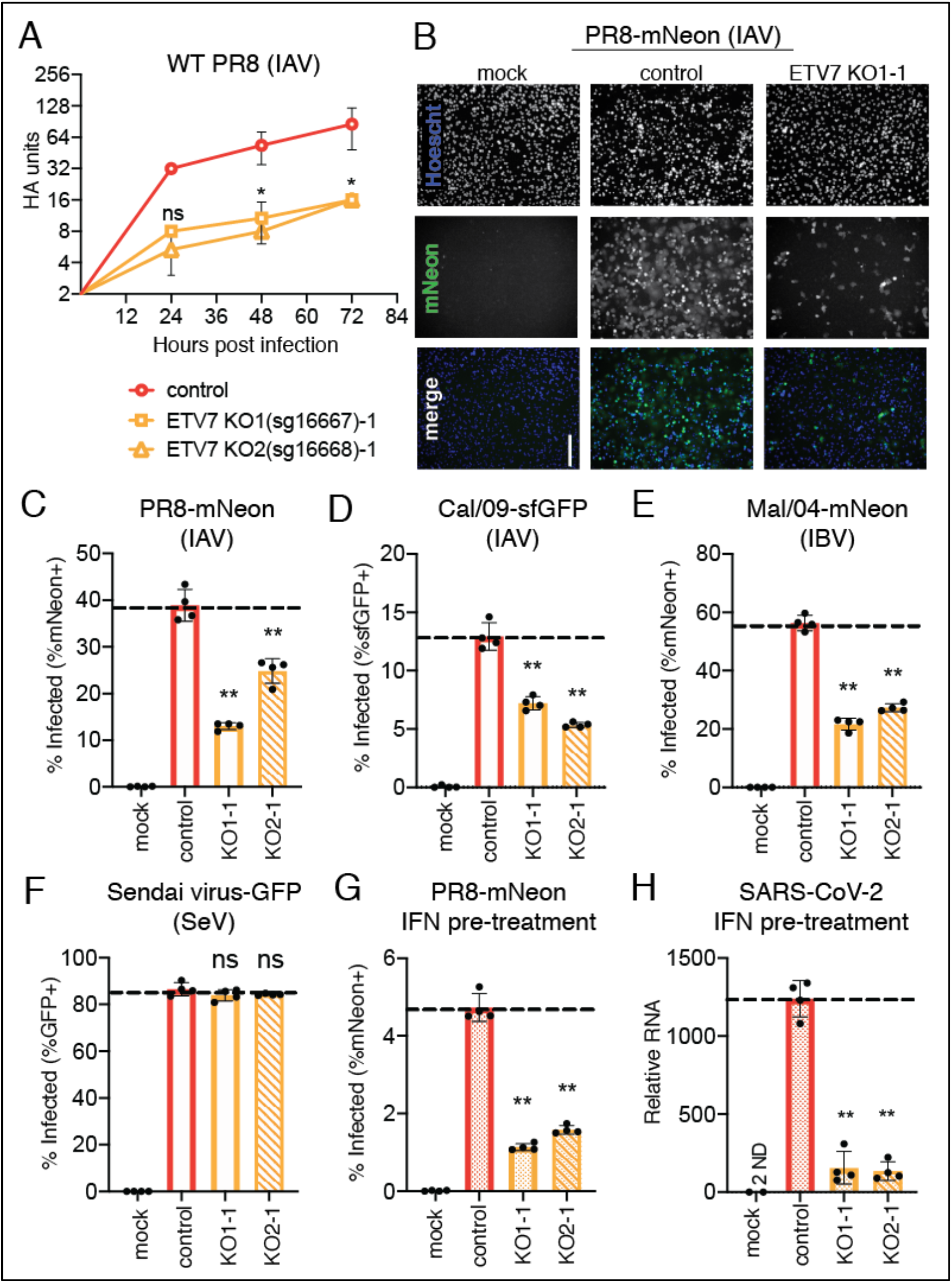
Loss of ETV7 leads to restricted growth of multiple strains of influenza virus and SARS-CoV-2 with IFN treatment. (**A**) Hemagglutination (HA) assay of virus collected at indicated time points from non-targeting control and ETV7 KO A549 clonal cell lines after infection with WT PR8 virus (MOI=0.05, multicycle infection) (data shown as mean ± SD, n=3). (**B**) Control and ETV7 KO A549 clonal cell lines after mock or PR8-mNeon reporter virus infection (24 h, MOI=0.1, multicycle infection). Green = mNeon, blue = nuclei. Scale bar, 200 μm. (**C**) Flow cytometry quantification of control and ETV7 KO A549 clonal cell lines after infection with PR8-mNeon reporter virus (24 h, MOI=0.1, multicycle infection) (data shown as mean ± SD, n=4). (**D, E**) Percentage of infected (reporter+) cells from ETV7 KO A549 clonal cell lines compared to a control cell line after infection with (**D**) Cal/09-sfGFP or (**E**) Mal/04-mNeon reporter viruses (24 h, multicycle infection) (data shown as mean ± SD, n=4). (**F**) Percentage of infected (reporter+) cells from ETV7 KO A549 clonal cell lines compared to a control cell line after infection with Sendai-GFP reporter virus (24 h) (data shown as mean ± SD, n=4). (**G**) Percentage of infected (reporter+) cells from ETV7 KO A549 clonal cell lines compared to a control cell line after pre-treatment with IFN-α (100 U/ml, 6 h) then infection with PR8-mNeon reporter virus (24 h, MOI=0.1, multicycle) (data shown as mean ± SD, n=4). (**H**) qPCR detecting SARS-CoV-2 N vRNA in infected cells from ETV7 KO A549 clonal cell lines compared to a control cell line after pre-treatment with IFN-α (1000 U/ml, 6 h) then infection with SARS-CoV-2 virus (24 h, MOI=0.1) (data shown as mean ± SD, n=4). For all panels: P-values calculated using unpaired, two-tailed Student’s t-tests (*p<0.05, **p<0.001) compared to infected, non-targeting sgRNA control samples.

In addition to physiological induction of IFN after infection, exogenous IFN treatment is frequently utilized or proposed as an antiviral therapy (*7*), either when there is difficulty developing targeted antivirals (e.g. hepatitis C virus (*54*)) or there is an unexpected emergence of a viral pathogen (e.g. swine or highly pathogenic avian influenza viruses (*55*), SARS-CoV (*56*), Ebola virus (*57*), SARS-CoV-2 (*58*–*60*)). Therefore, we wanted to assess whether suppressing ETV7 could be an effective strategy to enhance the antiviral effects of therapeutic IFN administration. In support of this idea, we found treatment of ETV7 KO cells with IFN-αprior to infection with PR8-mNeon led to an additional 5-fold enhancement of viral restriction compared to IFN-treated WT control cells (**Fig. 4G**). Finally, since recent reports indicate that although SARS-CoV-2 infections fail to induce a strong IFN responses (*61*–*63*), the virus is susceptible to IFN treatment (*64*–*66*), we wanted to test if loss of ETV7 would potentiate IFN-mediated suppression of SARS-CoV-2. With the same experimental design as the influenza virus experiment, we found that pre-treatment of ETV7 KO cells with IFN-α resulted in a further 10-fold reduction in SARS-CoV-2 vRNA compared to IFN-treated control cells (**Fig. 4H**). This enhanced control was likely mediated via the ISG LY6E, which potently restricts SARS-CoV-2 (*67*, *68*) and we found to be regulated by ETV7 (**Fig. S4B**). Together, these experiments demonstrate the potential of targeting ETV7 to enhance either the physiological or therapeutic effects of IFN to control viral infection.

## Discussion

In this study, we performed a CRISPR activation screen to identify negative regulators of the type I IFN response. Specifically, we were interested in negative regulators that contribute to the types of differentiated ISG profiles that are essential for effective control of viral infections. From this screen, we identified ETV7 as a negative IFN regulator and subsequently showed it acts as a transcription factor to repress subsets of ISGs through its recognized DNA binding motif. We also demonstrated that the regulatory activity of ETV7 impacts the replication and spread of multiple strains of influenza viruses. ETV7 loss also increased the antiviral effects of exogenous interferon treatment against an H1N1 influenza virus and SARS-CoV-2. These findings demonstrate the importance of ETV7 in fine-tuning the IFN response through specificity and transcriptional repression to regulate antiviral ISG targets, and its potential as a target to enhance antiviral IFN-based therapeutics.

ETV7 is a member of the ETS family of transcription factors. This family performs diverse functions despite recognizing the same core DNA sequence, GGAA. Because these factors share the same core motif, specificities are gained in other ways such as expression patterns (cell type, basal expression, or immune pathway induction), active or repressive activity, and potential binding partners (*69*, *70*). ETV7 is induced as an ISG and is repressive, whereas most ETS transcription factors are activators, including ETS transcription factors recognized to assist in the induction of ISG expression (e.g. ELF1 (*25*), PU.1 (*71*)). We also found repeatedly in our work that the reporters and genes impacted by ETV7 contained ETS sites within ISRE sequences (**Fig. 2E-H, S2G-I, S4B**). This is not unexpected because it is recognized that many ETS family members bind extended motifs similar to ISREs called ETS-IRF composite elements (EICE) (*72*). EICE-associated activity is reported to require an IRF binding partner to direct ETS transcription factor activity (*71*, *73*); therefore, it is likely ETV7 has an IRF binding partner. If ETV7 does require a binding partner, this protein’s induction and distribution likely contribute to the timing, gene targets, and activity of ETV7 during the IFN response. It is known that IRFs can be basally expressed (IRF2, IRF3), IFN-induced (IRF1, IRF7), or IFN-responsive (IRF9) (*72*), and the availability of a binding partner could dramatically affect the timing and magnitude of effects on EICE-controlled ISGs. Future work will define if ETV7 has specific binding partners and how those interactions may contribute to the nonuniform, repressive activity of ETV7 during the type I IFN response reported in this study.

IFN-induced regulators control the magnitude and duration of IFN responses in addition to the temporal regulation of specific waves of ISGs (*74*). These coordinated waves of ISG induction can peak early or late during the IFN response and are thought to correspond to specific stages of virus replication or immune processes (*1*, *6*). We compared the induction of ETV7 with IFI44L, IFIT1, ISG15 and RSAD2, and we observed ETV7 is upregulated at earlier time points than these prototypical ISGs (**Fig. 3A**). We expanded our analysis to published datasets of human gene expression during respiratory infections and concluded that ETV7 is generally induced earlier than many ISGs (*75*). Although not the focus of our study, ETV7’s early induction pattern suggests it may be a key regulator of the first stages of IFN-mediated gene induction. We favor a model wherein early ETV7 expression is responsible for reducing the accumulation, or delaying the expression, of ISGs controlled by ISREs and promoters containing ETS motifs.

ETV7 is induced during infections across many vertebrate species (*76*, *77*), indicating a potential conserved, relevant role in the immune response; however, ETV7 has been lost in mice and closely related rodents (*78*). Since mice and rodents have an intact interferon response pathway, a natural question is: how are the activities of ETV7 being accounted for in these animals? While we have no clear answer from the data in this study, it is well-recognized that IFN responses contain many redundancies (*41*). Accordingly, we believe other ETS family members, potentially the closely related ETV6 (which is also induced by IFNs), may perform the role of ETV7 in mice (*79*). Future studies will be required to test the hypothesis that mice induce an ETV7-related alternative during the type I IFN response.

Another important question is why the IFN-induced activity of ETV7 has been maintained throughout evolution. In this report, we provide evidence that ETV7’s activity reduces a cell’s ability to restrict virus infection; this seems counterintuitive to ETV7 benefitting the host. We hypothesize that regulators like ETV7 are important to prevent excessive inflammatory signaling. It is appreciated that negative regulators of the IFN response are required to prevent extreme and prolonged immune responses, which are associated with poor disease outcomes after infection (*80*–*82*). ETV7 potentially contributes to the cumulative activities of negative IFN regulators to limit IFN responses during pathogen clearance. Additionally, it stands to reason that individual ISGs have different toxic effects on the cell. It is tempting to speculate that ETV7 suppresses ISGs whose accumulation is particularly harmful to cell viability and host recovery after infection.

Additionally, the relevance of controlled IFN responses goes beyond infectious disease; patients with dysfunctional USP18, a negative regulator of the IFN response, develop a type I interferonopathy that results in a severe pseudo-TORCH (Toxoplasmosis, Other agents, Rubella, Cytomegalovirus, and Herpes simplex) syndrome (*83*). Mouse knockouts for other negative regulators of the IFN response (SOCS1, SOCS3, USP18) also develop non-pathogen associated, chronic inflammatory diseases (*84*–*87*). ETV7’s lack of a murine homolog eliminates an easily generated animal knockout model to experimentally show ETV7’s relevance as a general innate immune repressor. However, genome-wide association studies (GWAS) have linked ETV7 to autoimmune diseases including rheumatoid arthritis and multiple sclerosis (*88*, *89*); both of these autoimmune diseases have evidence of enhanced ISG expression (*90*, *91*). Thus, although the specific contributions of ETV7 activity to IFN regulation are currently undefined, its potential role is not limited to viral infections.

In conclusion, this study identified ETV7 as a negative regulator of the type I IFN response. Previously, ETV7 was appreciated to be an ISG; however, a specific function during the IFN response was unknown. We determined that ETV7 acts as a transcription factor to target specific ISGs for repression, potentially contributing to the complex ISG transcriptional landscape. Additionally, many of the ETV7-regulated ISGs restrict respiratory viruses (*42*, *67*, *68*), and we showed that loss of ETV7 can further enhance the ability of type I IFN to control influenza virus and SARS-CoV-2 replication. Further work is required to understand the complexity of IFN regulation, while therapeutic targeting of factors like ETV7 could lead to the development of a new class of host-directed antivirals that can enhance or tailor ISG responses to specific viruses.

## Materials and Methods

### Cloning

To generate reporters sensitive to IFN, we designed gBlocks (IDT) containing ISREs to be cloned into the pTRIP vector ahead of a minimal CMV promoter controlling expression of sfGFP. To clone and express the open reading frames (ORFs) of our screen hits, we designed primers for cloning into the pLEX-MCS vector using Gibson Assembly (NEB). To amplify ETV7, NUP153 and USP18 we used cDNA templates from Transomic Technologies. To amplify C1GALT1 and EIF2AK1 we used RNA from IFN-stimulated A549 cells. The DNA binding mutant ETV7, ETV7(KALK) (*35*), and codon-optimized SOCS1 expression plasmids were generated using a gBlock (IDT). FLAG-tagged ETV7, ETV7(FLAG), was generated using a primer including the FLAG tag. Non-targeting and ETV7-targeting CRISPR KO sgRNAs were cloned by annealing oligos encoding the desired sgRNA sequence and ligating them directly into the lentiCRISPRv2 vector (Addgene). DNA was transformed into NEB 5-alpha high efficiency competent cells. Insert size was verified with PCR and purified plasmids were sequenced using Sanger sequencing.

### Cells

All cells were obtained from ATCC and grown at 37°C in 5% CO_2_. A549 and 293T cells were grown in Dulbecco’s Modified Eagle Medium (DMEM) supplemented with 5% fetal bovine serum, GlutaMAX, and penicillin-streptomycin. Madin-Darby canine kidney (MDCK) cells were grown in minimal essential media (MEM) supplemented with 5% fetal bovine serum, HEPES, NaHCO_3_, GlutaMAX, and penicillin-streptomycin. The A549 CRISPR-SAM cells were previously validated (*92*) and transduced with the IFN response reporter three times before being clonally selected. The A549 CRISPR KO cells were transduced (ETV7 KO1 = sg16667, ETV7 KO2 = sg16668, nontargeting control) and then selected using puromycin (10 μg/mL). A549-IFN response reporter ETV7 KO lines were selected with puromycin and used as pooled lines. A549 ETV7 KO lines were selected with puromycin and subsequently plated at a dilution to isolate single cells, which were grown until colonies of an appropriate size allowed for collection. These clonally selected colonies were grown up and verified to be KO lines (control, KO1-1, KO1-2, KO2-1, KO2-2) by sequencing the DNA lesions generated as a result of Cas9 endonuclease activity, results shown below. Guide sequences = underlined, insertions = bolded, deletions = dashes, exonic sequences = uppercase, intronic sequences = lowercase.

ETV7 KO guide 1 (sg16667)

WT 5’ CTG CCA TGC ACC G<u>CG GAG CAC GGG TTC GAG ATG</u> AAC GGA CGC GCC 3’ protein length = 342aa
KO1-1 allele 1 5’ CTG CCA TGC ACC GCG GA**A** GCA CGG GTT CGA GAT GAA CGG ACG CGC 3’ predicted protein length = 104aa
KO1-1 allele 2 5’ CTG CCA TGC ACC GCG GA**A** GCA CGG GTT CGA GAT GAA CGG ACG CGC 3’ predicted protein length = 104aa
KO1-2 allele 1 5’ CTG CCA TGC ACC GCG GA**A** GCA CGG GTT CGA GAT GAA CGG ACG CGC 3’ predicted protein length = 104aa
KO1-2 allele 2 5’ CTG CCA TGC ACC GCG GA**A** GCA CGG GTT CGA GAT GAA CGG ACG CGC 3’ predicted protein length = 104aa

ETV7 KO guide 2 (sg16668)

WT 5’ GCG ATG CCG CAG GCC <u>CCC ATT GAC GGC AGG ATC GC</u>T Ggtgagtgggagg 3’ protein length = 342aa
KO2-1 allele 1 5’ GCG ATG CCG CAG GCC CCC ATT GAC G-- --- --- -CT Ggtgagtgggagg 3’ predicted protein length = 333aa
KO2-1 allele 2 5’ GCG ATG CCG CAG GCC CCC ATT GAC GGC AGG --- --- --tgagtgggagg 3’ predicted protein length = unknown, loss of splice site
KO2-2 allele 1 5’ GCG ATG CCG CAG GCC CCC ATT GAC G-- --- --- -CT Ggtgagtgggagg 3’ predicted protein length = 333aa
KO2-2 allele 2 5’ GCG ATG CCG CAG GCC CCC ATT GAC GGC AGG -T**GA** CGC TGgtgagtgggagg 3’ predicted protein length = 220aa

### Flow Cytometry

Cells were trypsinized and analyzed on a Fortessa X-20 (BD) machine with standard laser and filter combinations. Data was visualized and processed with FlowJo software.

### CRISPR Activation Screen

The sgRNA library was packaged into lentivirus as previously described (*92*). After packaging and titering the lentivirus, 2×10^8^ A549-CRISPR-SAM-IFN response reporter cells were seeded onto 15 cm plates (10 plates total). The next day they were transduced with the packaged sgRNA library (MOI=0.5). After 48 h, the transduced cells were split and half were collected as a transduction control, while the remaining half were plated back onto 15 cm plates. The next day, cells were treated with IFN-α(4×10^3^ U/mL) for 6 h. Cells were then collected and sorted on a Beckman Coulter Astrios cell sorter. Specifically, gates were set to sort GFP-negative cells as the population of interest, as well as GFP-positive cells as a control population of cells still capable of signaling. This screen was performed in duplicate. Genomic DNA was extracted from sorted cells using the Zymo Quick gDNA micro prep kit. PCR was subsequently performed using barcoded primers as previously described using the NEB Next High Fidelity 2x PCR master mix (*92*). PCR bands were gel purified using the Thermo GeneJET gel extraction kit. Samples were then sequenced via next-generation Illumina MiSeq using paired-end 150 bp reads.

### Screen Analysis

Raw MiSeq read files were aligned to the CRISPR SAM sgRNA library and raw reads for each sgRNA were counted using the MAGeCK pipeline (*29*). sgRNA enrichment was determined using the generated count files and the MAGeCK-MLE analysis pipeline. Genes were sorted based on z-score and determined to be significantly enriched if their z-score was at least two standard deviations above the average z-score of the entire sorted population.

### Western Blotting

Cells were trypsinized and 1×10^6^ cells were pelleted at 800 x g for 5 min. Equal amounts of cellular material were loaded into 4-20% acrylamide gels (Bio-Rad) and imaged using a ChemiDoc Imaging System (Bio-Rad). Protein was transferred to a nitrocellulose membrane at 60V for 60 min. PBS with 5% (w/v) non-fat dried milk and 0.1% Tween-20 were used to block for 1 h at 4°C. Primary antibodies were then incubated with the membrane overnight at 4°C. Antibodies used were rabbit anti-ETV7 (Sigma, HPA029033), rabbit anti-IFIT1 (Cell Signaling, D2X9Z), mouse anti-FLAG (Sigma, F3165) and mouse anti-αtubulin (Sigma, T5168). Membranes were washed five times in PBS with 0.1% Tween-20 and then an anti-rabbit-HRP (Thermo, A16104) or anti-mouse-HRP (Thermo, A16072) secondary antibody was added for 1 h. The membrane was then washed five times and Clarity or Clarity Max ECL substrate (Bio-Rad) was added before being exposed to film and developed.

### RT-qPCR

For all experiments except SARS-CoV-2 infection experiments, total RNA was collected and prepared using Monarch Total RNA Miniprep Kits (NEB). For SARS-CoV-2 experiments, cells were collected in TRIzol (Invitrogen) and RNA was isolated using Phasemaker tubes (Invitrogen). One-step RT-qPCR was performed with commercial TaqMan assays from Thermo for ETV7 (Hs00903229_m1), C1GALT1 (Hs00863329_g1), NUP153 (Hs01018919_m1), ISG15 (Hs00196051_m1), MX1 (Hs00895608_m1), IFIT1 (Hs00356631_g1), IFI44L (Hs00915292_m1), RSAD2 (Hs00895608_m1), IFITM1 (Hs00705137_s1), OAS1 (Hs00973635_m1), IFIT2 (Hs00533665_m1), IFIT3 (Hs01922752_s1), IFI44 (Hs00197427_m1), BST2 (Hs01561315_m1), and ACTB (Hs01060665_g1) and primers/probe from BEI targeting the SARS-CoV-2 N region using the EXPRESS One-Step Superscript qRT-PCR Kit on an Applied Biosystems StepOnePlus or QuantStudio 3 instrument. RNA was normalized using an endogenous 18S rRNA primer/probe set (Applied Biosystems).

### DNA pull-down assay

Nuclear extract from 293T cells expressing ETV7 and treated with IFN-α (100 U/mL, 9 h) was generated using NE-PER extraction reagents (Thermo Scientific). This nuclear extract was incubated with poly(dI-dC) as nonspecific competitor DNA, 10x ligase buffer (Invitrogen), 200 mM EDTA, biotinylated DNA bait (either WT or −ETS ISG15 ISRE (*93*)), and 20x excess nonbiotinylated DNA competitor for 30 min at room temperature. This mixture was then incubated with streptavidin-coated magnetic beads (Invitrogen), followed by four washes with TTBS. Proteins bound to the biotinylated DNA bait captured by the streptavidin beads were eluted with protein sample buffer and then detected by Western blot. Input (nuclear extract) was diluted 1:200 for loading. No bait (control) contained no biotinylated DNA.

### RNA sequencing

293T cells were transfected with ETV7- or control-expressing plasmids and selected using puromycin (20 μg/mL) for 24 h before treatment with IFN-α (100 U/mL). Total RNA was collected at 9 h post-IFN treatment using Monarch Total RNA Miniprep Kits (NEB). RNA was prepped for RNA sequencing submission using the NEBNext Poly(A) mRNA Magnetic Isolation Module (NEB), NEBNext Ultra II RNA Library Prep Kit for Illumina (NEB), and NEBNext Multiplex Oligos for Illumina (NEB). Samples were analyzed on one lane of an Illumina HiSeq 4000 using 50 bp single strand reads. Mapping of the raw reads to the human hg19 reference genome was accomplished using a custom application on the Illumina BaseSpace Sequence Hub (*94*). After data normalization, average read values were compared across samples. For comparisons in which some samples had zero reads detected for a specific gene, one read was added to all values in the sample to complete analyses that required non-zero values. The heat map shows genes upregulated 2.5-fold (after normalization) with IFN treatment in the control samples. Values shown are z-scores.

### ChIP-qPCR

293T cells were transfected with a FLAG-tagged ETV7-expressing plasmid, ETV7(FLAG), 48 h before treatment with IFN-α (100 U/mL, 9 h). Chromatin immunoprecipitation was performed using the ChIP-IT Express Enzymatic kit according to the manufacturer’s directions (Active Motif). DNA was enzymatically sheared for 10 min. ChIP antibodies included mouse IgG (Active Motif), rabbit anti-FLAG (Cell Signaling, 14793S), and mouse anti-FLAG (Sigma, F3165). Quantitative PCR was performed using SYBR Green (Bio-Rad) and primers targeting the Chr12 gene desert (Active Motif) or IFI44L promoter (forward primer = 5’ TTTCATGCCTGCCTACATAC 3’, reverse primer = 5’ ATGCCAACTGCCACTAAC 3’) in a region containing two potential ISRE sequences overlapping with ETS sites, and analyzed using the ChIP-IT qPCR Analysis kit (Active Motif).

### Viruses

PR8-mNeon was generated via insertion of the mNeon fluorescent gene (*95*) into segment 4 of the virus (*48*). Mal/04-mNeon was generated by inserting the mNeon fluorescent gene (*95*) into segment 4 of the Mal/04 genome (*49*). Cal/09-sfGFP was generated via insertion of the sfGFP gene (*96*) into segment 8 of the virus using the same scheme previously used to insert Cre recombinase (*97*). Sendai-GFP was a gift from Benhur Lee (*98*). For influenza virus infections, cells were either mock- or virus-infected for 1 h and then cultured in OptiMEM supplemented with bovine serum albumin (BSA), penicillin-streptomycin, and 0.2 μg/mL TPCK-treated trypsin protease (Sigma). For Sendai infections, cells were infected for 1 h and then cultured in DMEM. For SARS-CoV-2 infections, cells were washed with PBS before infection with SARS-CoV-2 isolate USA-WA1/2020 from BEI Resources in 2% FBS DMEM infection media for 1 hour. PR8 WT, PR8-mNeon, Cal/09-sfGFP, Sendai-GFP, and SARS-CoV-2 were incubated at 37°C, Mal/04-mNeon was incubated at 33°C.

### Viral Growth Assays

Hemagglutination (HA) assays to measure the number of viral particles were performed by diluting influenza infected cell supernatants collected at the indicated time points in cold PBS. An equal amount of chicken blood diluted 1:40 in PBS was mixed with serially diluted virus and incubated at 4°C for 2-3 h before scoring. Infectious viral titers were determined using standard plaque assay procedures on MDCK cells. Infected cell supernatants were collected at 18 h, serially diluted, and used to infect confluent 6-well plates for 1 h before removing the virus and adding the agar overlay. Cells were then incubated at 37°C for 48 h before being fixed in 4% PFA overnight. The 4% PFA was then aspirated, and the agar layer was removed before washing cells with PBS. Serum from WT PR8 infected mice was diluted 1:2,000 in antibody dilution buffer (5% (w/v) non-fat dried milk and 0.05% Tween-20 in PBS) and incubated on cells at 4°C overnight. Cells were then washed twice with PBS and incubated for 1 h with anti-mouse IgG horseradish peroxidase (HRP)-conjugated sheep antibody (GE Healthcare) diluted 1:4,000 in antibody dilution buffer. Assays were then washed twice with PBS and exposed to 0.5 mL of TrueBlue peroxidase substrate (KPL) for 20 min. Plates were then washed with water and dried before plaques were counted.

## Supplementary Materials

Fig. S1. CRISPR activation screen hit identification and validation.

Fig. S2. ETV7 represses IFN-stimulated expression and directly binds an ISG promoter.

Fig. S3. ETV7 loss increases IFN-stimulated expression.

Fig. S4. ETV7 loss results in decreased viral titers and increased antiviral gene expression.

Table S1. Hits from CRISPRa screen for negative regulators of the type I IFN response.

Table S2. ISRE sequences identified in the literature.

Table S3. Gene set enrichment analysis of 200 most downregulated genes in ETV7-expressing cells without IFN treatment.

Table S4. Gene set enrichment analysis of 200 most downregulated genes in ETV7-expressing cells after 9hr IFN treatment.

Data file S1. CRISPRa screen for negative regulators of the type I IFN response sgRNA sequences, transduction read counts, and screen read counts.

Data file S2. CRISPRa screen for negative regulators of the type I IFN response MAGeCK z-scores and potential hit cutoffs.

Data file S3. RNA sequencing with ETV7 overexpression and IFN treatment reads and results.

## Acknowledgements

We acknowledge assistance from Mike Cook and the Duke Cancer Institute Flow Cytometry Core. We thank Robert Lefkowitz and his laboratory for assistance with and use of the ChemiDoc Imaging System. We would like to thank Dr. Clare Smith for help establishing SARS-CoV-2 viral infection assays at BSL3. We also thank Ephraim Tsalik, Micah McClain, and Bandita Gershon for helpful discussions and Ben Chambers, Stacy Webb, and other members of the Heaton lab for critical reading of the manuscript.

## Author contributions

H.M.F., A.T.H. and N.S.H. designed the study. H.M.F., A.T.H., and B.E.H. performed experiments. H.M.F., A.T.H. and N.S.H. analyzed data. H.M.F. and N.S.H. wrote the paper.

## Data and materials availability

All next generation sequencing data are available at NCBI GEO under accession number GSE140718. The following reagent was deposited by the Centers for Disease Control and Prevention and obtained through BEI Resources, NIAID, NIH: SARS-Related Coronavirus 2, Isolate USA-WA1/2020, NR-52281.

