## Supplemental Figures and Tables for "ETV7 limits antiviral gene expression and control of SARS-CoV-2 and influenza viruses"

**This file includes:**

Fig. S1. CRISPR activation screen hit identification and validation.
Fig. S2. ETV7 represses IFN-stimulated expression and directly binds an ISG promoter.
Fig. S3. ETV7 loss increases IFN-stimulated expression.
Fig. S4. ETV7 loss results in decreased viral titers and increased antiviral gene expression.
Table S1. Hits from CRISPRa screen for negative regulators of the type I IFN response.
Table S2. ISRE sequences identified in the literature.
Table S3. Gene set enrichment analysis of 200 most downregulated genes in ETV7-expressing cells without IFN treatment.
Table S4. Gene set enrichment analysis of 200 most downregulated genes in ETV7-expressing cells after 9hr IFN treatment.

**Other Supplementary Material for this manuscript includes the following:**

Data file S1. CRISPRa screen for negative regulators of the type I IFN response sgRNA sequences, transduction read counts, and screen read counts.
Data file S2. CRISPRa screen for negative regulators of the type I IFN response MAGeCK z-scores and potential hit cutoffs.
Data file S3. RNA sequencing with ETV7 overexpression and IFN treatment reads and results.

**Figure S1**

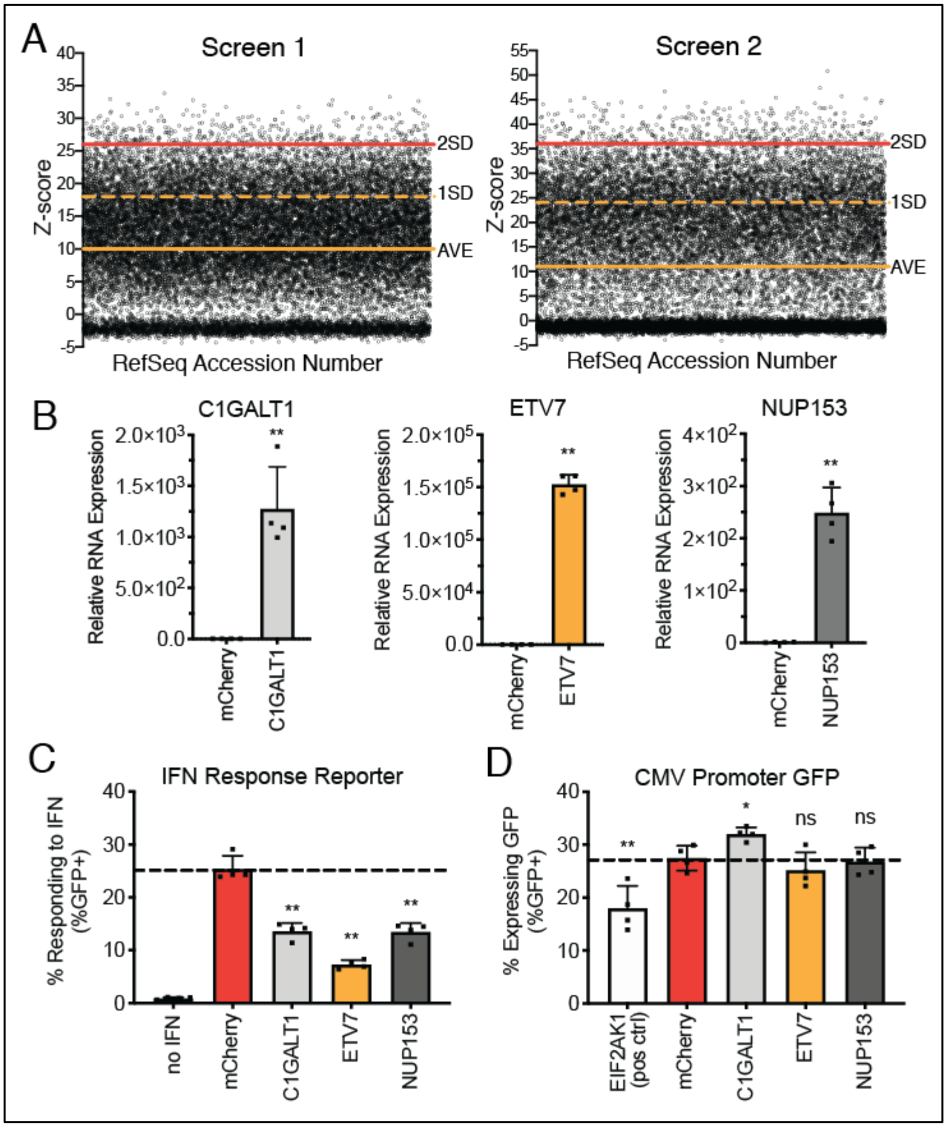


**Fig. S1. CRISPR activation screen hit identification and validation.** (**A**) Results of the two independent CRISPRa screens. Z-score values from the replicate screens with a cutoff of two SD from the mean (Screen 1 = 284 genes, Screen 2 = 392 genes) were used to identify top "hits". **(B)** RT-qPCR of three screen hits, C1GALT1, NUP153, and ETV7, in 293T cells confirms their overexpression after transfection of a pLex vector containing the respective gene. (**C**) Percentage of cells expressing GFP from the IFN response reporter with overexpression of the indicated genes compared to the mCherry-expressing control (data shown as mean ± SD, n=4). (**D**) Percentage of cells expressing GFP from a constitutively expressing plasmid in cells overexpressing the indicated genes (positive control = EIF2AK1/HRI, shuts off translation) compared to control (data shown as mean ± SD, n=4). For all panels: P-values calculated using unpaired, two-tailed Student's t-tests (*$p<0.05$, **$p<0.001$) compared to mCherry-expressing control samples.

**Figure S2**

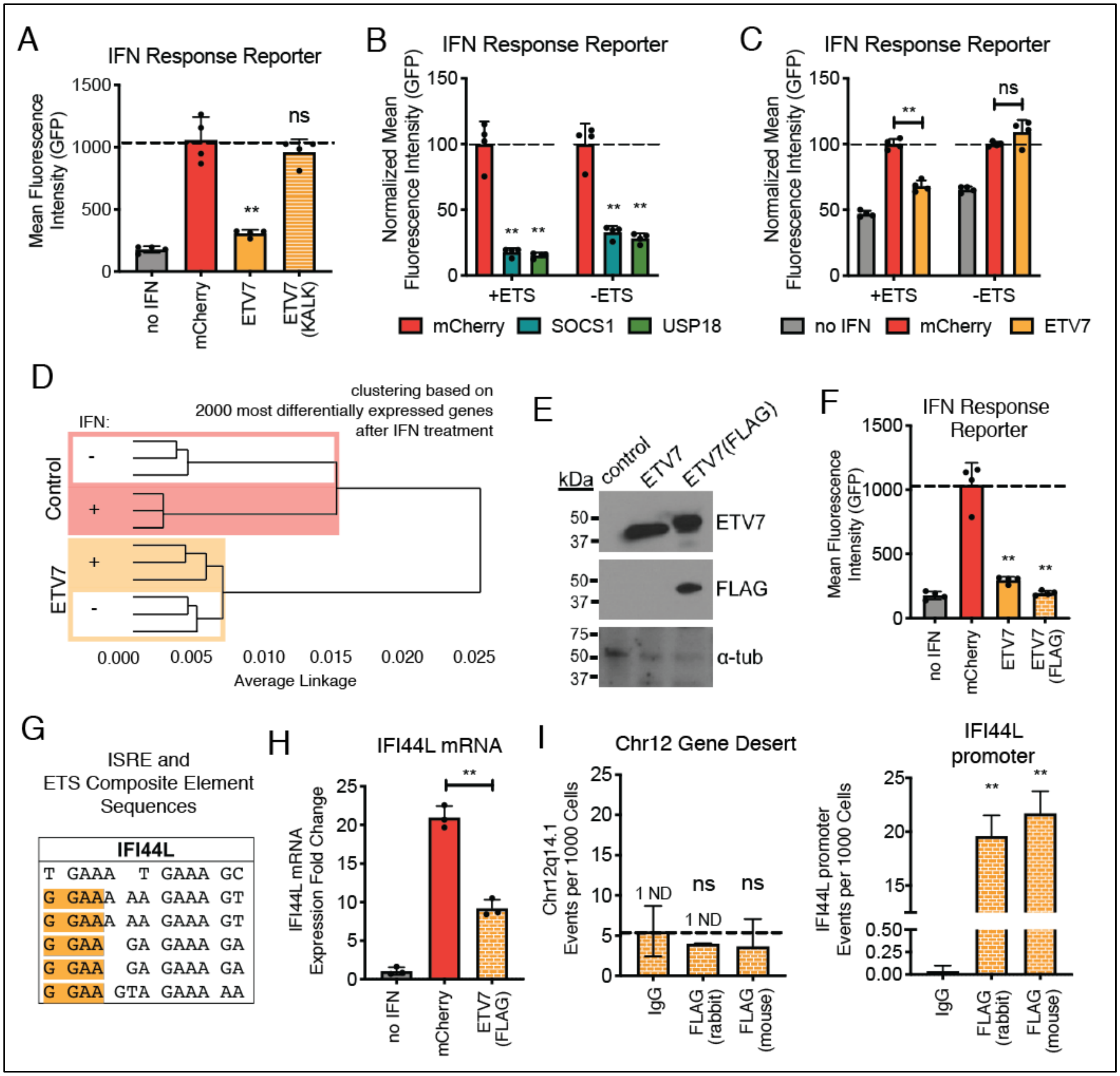


**Fig. S2. ETV7 represses IFN-stimulated expression and directly binds an ISG promoter.** (**A**) Brightness of 293T cells expressing GFP from the IFN response reporter with overexpression of WT or DNA binding mutant (KALK) ETV7 after IFN-α treatment (100 U/mL, 9 h) compared to control (data shown as mean ± SD, n=4, statistical analysis relative to IFN-stimulated, mCherry-expressing control samples). (**B**) Brightness of 293T cells expressing GFP from IFN response reporters either containing or not containing ETS sites after overexpression of known negative regulators of the IFN response (SOCS1, USP18) and IFN-α treatment (100 U/mL, 9 h) compared to mCherry-expressing control (data shown as mean ± SD, n=4, statistical analysis relative to IFN-stimulated, mCherry-expressing control samples). (**C**) Normalized brightness of 293T cells expressing GFP from IFN response reporters either containing or not containing ETS sites after overexpression of ETV7 and IFN-α treatment (100 U/mL, 6 h) compared to mCherry-expressing control (data shown as mean ± SD, n=4, statistical analysis relative to IFN-stimulated, mCherry-expressing control samples). (**D**) Dendrogram of genes most differentially expressed in cells overexpressing either a control protein (mCherry) or ETV7 before and after IFN-α treatment (100 U/mL, 9 h) as measured using RNA sequencing. Three independent, biological replicates per condition. Red box highlights control samples, yellow box highlights ETV7-expressing samples, shading indicates IFN-stimulated samples. The box width indicates the linkage distance between samples before and after IFN, indicating control cells' transcriptional profile is more diverged after IFN treatment compared to ETV7-expressing cells. Generated using Heatmapper (*100*). (**E**) Western blot showing ETV7 protein and FLAG levels with overexpression of ETV7, FLAG-tagged ETV7, or control (mCherry) in 293T cells. α-tubulin was used to show loading. (**F**) Brightness of 293T cells expressing GFP from the IFN response reporter with overexpression of WT or FLAG-tagged ETV7 after IFN-α treatment (100 U/mL, 9 h) compared to control (data shown as mean ± SD, n=4, statistical analysis relative to IFN-stimulated, mCherry-expressing control samples). (**G**) Potential ISRE-like sequences in the IFI44L promoter. ETS sites (GGAA) highlighted in yellow. (**H**) IFI44L mRNA expression levels measured using RT-qPCR after IFN-α treatment (100 U/mL, 9 h) with FLAG-tagged ETV7 expression (data shown as mean ± SD, n=3, statistical analysis relative to IFN-stimulated, mCherry-expressing control samples). (**I**) ChIP-qPCR of the Chr12 gene desert (negative control) and the IFI44L promoter region in 293T cells expressing FLAG-tagged ETV7 and treated with IFN-α (100 U/ml, 9 h), immunoprecipitated using nonspecific (IgG) or targeted (anti-FLAG, rabbit or mouse) antibodies (data shown as mean ± SD, n=3, statistical analysis relative to IgG control samples). For all panels: P-values calculated using unpaired, two-tailed Student's t-tests (*p<0.05, **p<0.001) unless otherwise noted.

**Figure S3**

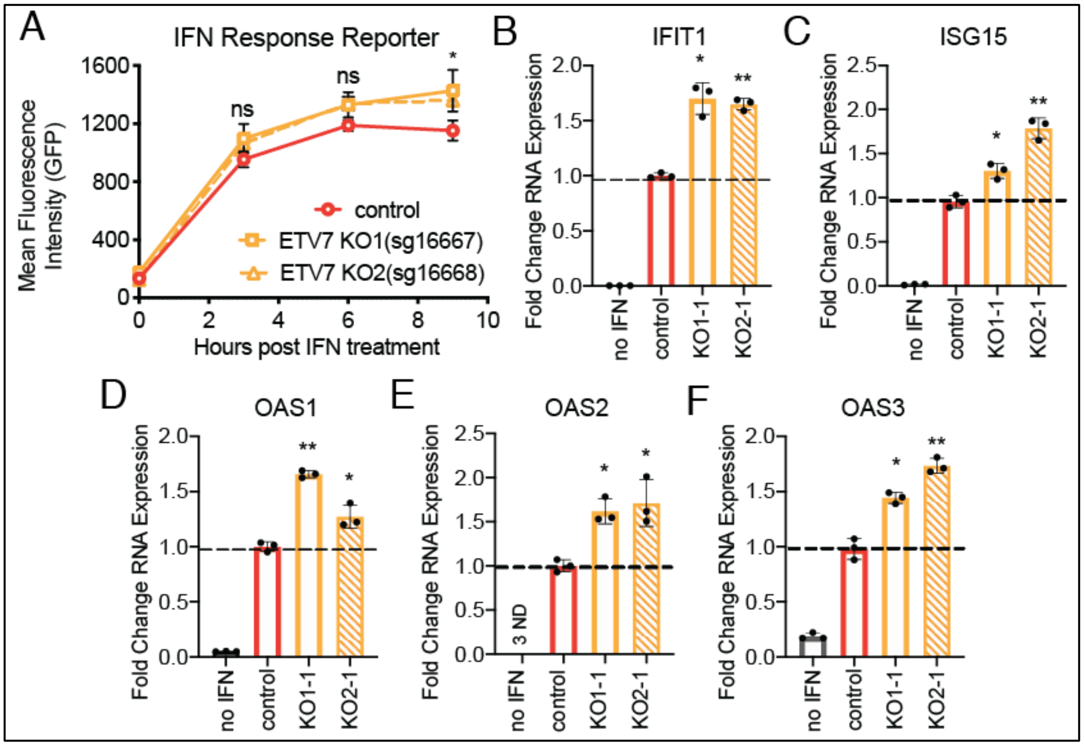


**Fig. S3. ETV7 loss increases IFN-stimulated expression.** (**A**) Brightness of cells expressing GFP from the IFN response reporter in A549 ETV7 KO cells (pooled, 2 different guides) after IFN-α treatment (1000 U/mL, 6 h) compared to non-targeting control cells (data shown as mean ± SD, n=3). (**B-F**) mRNA levels of ISGs in control and ETV7 KO A549 clonal cells after IFN-α treatment (100 U/mL, 9 h) (data shown as mean ± SD, n=3). For all panels: P-values calculated using unpaired, two-tailed Student's t-tests (*p<0.05, **p<0.001) compared to IFN-stimulated, non-targeting sgRNA control samples unless otherwise indicated.

**Figure S4**

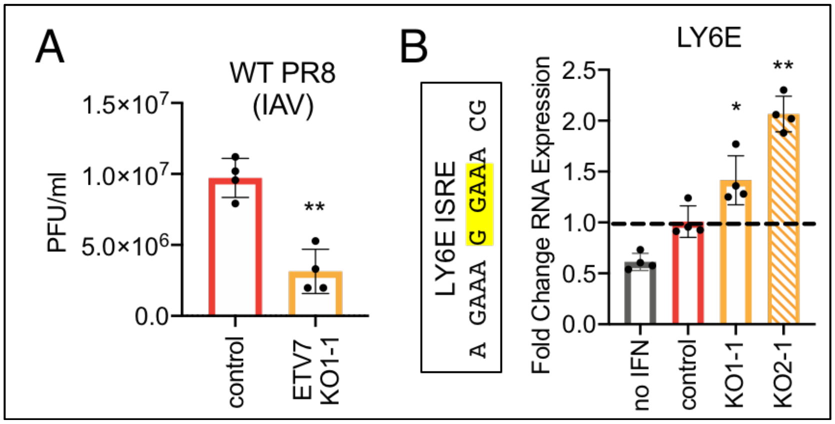


**Fig. S4. ETV7 loss results in decreased viral titers and increased antiviral gene expression.** (**A**) Titer of virus collected from control and ETV7 KO1(sg16667)-1 A549 cells after infection with WT PR8 virus (18 h, MOI=0.05, multicycle infection) (data shown as mean ± SD, n=4). (**B**) ISRE sequence in the LY6E promoter. ETS site (GGAA) highlighted in yellow. mRNA levels of LY6E in control and ETV7 KO A549 clonal cells after IFN-α treatment (100 U/mL, 9 h) (data shown as mean ± SD, n=4). For all panels: P-values calculated using unpaired, two-tailed Student's t-tests (*p<0.05, **p<0.001) compared to infected or IFN-stimulated, non-targeting sgRNA control samples.

**Table S1. Hits from CRISPRa screen for negative regulators of the type I IFN response.**

| **Gene** | **Induced by IFN >2x** (Interferome) | **HGNC Official Full Name** |
|---|---|---|
| C1GALT1 | Yes | core 1 synthase, glycoprotein-N-acetylgalactosamine 3-beta-galactosyltransferase 1 |
| CASC3 | No | CASC3 exon junction complex subunit |
| ETV7 | Yes | ETS variant transcription factor 7 |
| GGT1 | No | gamma-glutamyltransferase 1 |
| GOLGA6D | No | golgin A6 family member D |
| IMPAD1 | No | inositol monophosphatase domain containing 1 |
| LAPTM5 | No | lysosomal protein transmembrane 5 |
| NUP153 | Yes | nucleoporin 153 |
| PCP2 | No | Purkinje cell protein 2 |
| RRM2B | No | ribonucleotide reductase regulatory TP53 inducible subunit M2B |

Ten hits identified using overlap between two replicate screens. Hits selected for validation were determined to be induced at least two-fold after IFN treatment in the Interferome database (*30*).

**Table S2. ISRE sequences identified in the literature.**

| Gene | ISRE sequence | Reference | Ref # |
|---|---|---|---|
| *Consensus ISRE* | N GAAA NN GAAA CT | Bluyssen et al. (1994) | *101* |
| BST2 | G GAAA CT GAAA CT | Ohtomo et al. (1999) | *102* |
| IFI35 | G GAAA T GAAA GT | Yang et al. (2012) | *103* |
| IFIT1 | G GAAA GT GAAA CT | Bluyssen et al. (1994) | *101* |
| | G GAAA CC GAAA GG | Wathelet et al. (1998) | *104* |
| IFIT2 | G GAAA GT GAAA CT | Levy et al. (1988) | *10* |
| IRF9 | A GAA CT GAAA CT | Testoni et al. (2011) | *105* |
| ISG15 | G GAAA CC GAAA CT | Testoni et al. (2011) | *105* |
| | G GAAA GG GAAA CC | Testoni et al. (2011) | *105* |
| OAS1 | G GAAA C GAAA CC | Rutherford et al. (1988) | *106* |
| OAS2 | G GAAA CT GAAA CT | Wang and Floyd-Smith (1997) | *107* |
| OAS3 | G GAAA AC GAAA CC | Rebouillat et al. (2000) | *108* |
| | C GAAA CT GAAA GC | Rebouillat et al. (2000) | *108* |
| OASL | A GAA TC GAAA CT | Wang et al. (2010) | *109* |

Consensus ISRE with potential ETS sites highlighted in blue. ISGs and their identified ISRE sequences with ETS sites highlighted in yellow and non-ETS sites highlighted in gray.

**Table S3. Gene set enrichment analysis of 200 most downregulated genes in ETV7-expressing cells without IFN treatment.**

| Source | Term | p-value |
|---|---|---|
| TF | Factor: Fli-1; motif: ACCGGAAGYN | 5.11176E-06 |
| TF | Factor: Erm; motif: ACCGGAAGTN | 1.13194E-05 |
| TF | Factor: c-Ets-2; motif: NCCGGAAGTG | 1.26284E-05 |
| TF | Factor: SAP-1; motif: NNCCGGAAGTGN | 2.48889E-05 |
| TF | Factor: Erg; motif: ACCGGAAGTN | 3.27746E-05 |
| TF | Factor: ETV4; motif: ACCGGAAGTN | 7.51935E-05 |
| TF | Factor: c-ets-1; motif: ACCGGAWGTN | 9.58909E-05 |
| TF | Factor: FLI-1; motif: NAYTTCCGGT | 0.000135473 |
| TF | Factor: PEA3; motif: NACCGGAAGTN | 0.000146003 |
| TF | Factor: Elf-1; motif: ACTTCCGGG | 0.000191389 |
| TF | Factor: ETS1; motif: ACCGGAARYN | 0.000213448 |
| TF | Factor: PEA3; motif: RCCGGAAGYN | 0.000250052 |
| GO:CC | RNA polymerase I transcription factor complex | 0.000387723 |
| TF | Factor: Elk-1; motif: RCCGGAAGTGN | 0.000476268 |
| TF | Factor: ER71; motif: ACCGGAARYN | 0.00062605 |
| TF | Factor: Erg; motif: NACCGGAARTN | 0.00084079 |
| TF | Factor: PEA3; motif: NACCGGAAGTN | 0.00124071 |
| TF | Factor: Fli-1; motif: NACCGGAARTN | 0.00139851 |
| TF | Factor: c-Ets-1; motif: NNNRCCGGAWRYNNNN | 0.00163403 |
| TF | Factor: ELK1; motif: ACCGGAAGTN | 0.00194463 |

Top 20 results from gProfiler (*37*) analysis, organized by p-value. gProfiler maps genes from a provided list to databases, such as Gene Ontology, KEGG, Reactome, miRTarBase, TRANSFAC, Human Protein Atlas, CORUM, and Human Phenotype Ontology, and determines significantly enriched terms. The gene list submitted was generated by comparing the percent read of the "no IFN" mCherry and ETV7 RNA sequencing results using fold change and sorting for the 200 most downregulated genes in the ETV7 dataset. Yellow highlight = ETS site, blue highlight = potential ETS site.

<u>Key</u>
GO:MF = Gene Ontology: molecular function (gene ontology)
GO:CC = Gene Ontology: cellular component (gene ontology)
GO: BP = Gene Ontology: biological process (gene ontology)
KEGG = KEGG (pathways)
REAC = Reactome (pathways)
WP = WikiPathways (pathways)
TF = TRANSFAC (regulatory motifs)
MIRNA = miRTarBase (miRNA targets)
HPA = Human Protein Atlas (tissue specificity)
CORUM = CORUM (protein complexes)
HP = Human Phenotype Ontology (human disease)

**Table S4. Gene set enrichment analysis of 200 most downregulated genes in ETV7-expressing cells after 9hr IFN treatment.**

| Source | Term | p-value |
|---|---|---|
| GO:BP | defense response to virus | 1.43E-15 |
| GO:BP | response to virus | 1.44E-14 |
| GO:BP | type I interferon signaling pathway | 2.78E-14 |
| GO:BP | cellular response to type I interferon | 2.78E-14 |
| REAC | Interferon alpha/beta signaling | 2.96E-14 |
| GO:BP | response to type I interferon | 4.66E-14 |
| GO:BP | negative regulation of viral genome replication | 9.64E-13 |
| GO:BP | negative regulation of viral life cycle | 5.75E-12 |
| GO:BP | negative regulation of viral process | 8.81E-11 |
| REAC | Interferon Signaling | 6.27E-10 |
| GO:BP | regulation of viral genome replication | 8.49E-10 |
| TF | Factor: IRF-9; motif: NYGAAACYGAAACYN | 1.55E-09 |
| TF | Factor: IRF; motif: NNGAAANTGAAANN | 2.97E-09 |
| GO:BP | regulation of viral life cycle | 2.18E-08 |
| GO:BP | viral genome replication | 2.67E-08 |
| TF | Factor: IRF-1; motif: STTTCACTTTCNNT | 2.77E-08 |
| TF | Factor: IRF-8; motif: NCGAAACYGAAACYN | 4.77E-08 |
| TF | Factor: IRF-8; motif: NCGAAACCGAAACYN | 6.24E-08 |
| TF | Factor: IRF; motif: NNGAAANTGAAANN | 7.28E-08 |
| TF | Factor: IRF-1; motif: NNRAAANNGAAASN | 1.38E-07 |

Top 20 results from gProfiler (*37*) analysis, organized by p-value. gProfiler maps genes from a provided list to databases, such as Gene Ontology, KEGG, Reactome, miRTarBase, TRANSFAC, Human Protein Atlas, CORUM, and Human Phenotype Ontology, and determines significantly enriched terms. The gene list submitted was generated by comparing the percent read of the "9hr IFN" mCherry and ETV7 RNA sequencing results using fold change and sorting for the 200 most downregulated genes in the ETV7 dataset. Yellow highlight = ETS site, blue highlight = potential ETS site.

Key

GO:MF = Gene Ontology: molecular function (gene ontology)
GO:CC = Gene Ontology: cellular component (gene ontology)
GO: BP = Gene Ontology: biological process (gene ontology)
KEGG = KEGG (pathways)
REAC = Reactome (pathways)
WP = WikiPathways (pathways)
TF = TRANSFAC (regulatory motifs)
MIRNA = miRTarBase (miRNA targets)
HPA = Human Protein Atlas (tissue specificity)
CORUM = CORUM (protein complexes)
HP = Human Phenotype Ontology (human disease)

**Data file S1.** CRISPRa screen for negative regulators of the type I IFN response sgRNA sequences, transduction read counts, and screen read counts.

**Data file S2.** CRISPRa screen for negative regulators of the type I IFN response MAGeCK z-scores and potential hit cutoffs.

**Data file S3.** RNA sequencing with ETV7 overexpression and IFN treatment reads and results.
